# Public health in genetic spaces: a statistical framework to optimize cluster-based outbreak detection

**DOI:** 10.1101/639997

**Authors:** Connor Chato, Marcia L. Kalish, Art F. Y. Poon

## Abstract

Genetic clustering is a popular method for characterizing variation in transmission rates for rapidly-evolving viruses, and could potentially be used to detect outbreaks in ‘near real time’. However, the statistical properties of clustering are poorly understood in this context, and there are no objective guidelines for setting clustering criteria. Here we develop a new statistical framework to optimize a genetic clustering method based on the ability to forecast new cases. We analyzed the pairwise Tamura-Nei (TN93) genetic distances for anonymized HIV-1 subtype B *pol* sequences from Seattle (*n* = 1, 653) and Middle Tennessee, USA (*n* = 2, 779), and northern Alberta, Canada (*n* = 809). Under varying TN93 thresholds, we fit two models to the distributions of new cases relative to clusters of known cases: (1) a null model that assumes cluster growth is strictly proportional to cluster size, *i.e.*, no variation in transmission rates among individuals; and (2) a weighted model that incorporates individual-level covariates, such as recency of diagnosis. The optimal threshold maximizes the difference in information loss between models, where covariates are used most effectively. Optimal TN93 thresholds varied substantially between data sets, *e.g.*, 0.0104 in Alberta and 0.016 in Seattle and Tennessee, such that the optimum for one population will potentially mis-direct prevention efforts in another. The range of thresholds where the weighted model conferred greater predictive accuracy tended to be narrow (*±*0.005 units), but the optimal threshold for a given population also tended to be stable over time. We also extended our method to demonstrate that variation in recency of HIV diagnosis among clusters was significantly more predictive of new cases than sample collection dates (ΔAIC*>* 50). These results demonstrate that one cannot rely on historical precedence or convention to configure genetic clustering methods for public health applications. Our framework not only provides an objective procedure to optimize a clustering method, but can also be used for variable selection in forecasting new cases.

## Background

Spatiotemporal clustering is a fundamental public health methodology for the detection of infectious disease outbreaks [1]. The colocalization of cases in space and time can reveal the existence of a common source, and cases within a cluster tend to be related by recent transmission events. For example, an automated space-time clustering method [2] was demonstrated to retrospectively detect outbreaks of nosocomial bacterial infection in a US-based hospital, including the outbreaks that were detected contemporaneously by the hospital’s pre-existing infection control program [3]. At a broader spatial scale, the same clustering method was recently used to identify outbreaks of severe acute respiratory infections over a five year period, using case data from a network of hospitals in Uganda [4]. Early detection of a cluster represents a potential opportunity for a targeted public health response to prevent additional cases. Space-time clustering may be less effective, however, for pathogens that can establish a chronic infection with a long asymptomatic period (*e.g.*, *Mycobacterium tuberculosis*, hepatitis C virus, or human immunodeficiency virus type 1; HIV-1) where the transmission event may have occurred months or years before diagnosis. Furthermore, pathogens with a relatively low per-act transmission rate present difficulties for space-time clustering because a single exposure in a specific location is unlikely to result in transmission. Under these circumstances, the spread of an epidemic is more likely to be shaped by a social network of repeated contacts between individuals, rather than shared venues.

For many infectious diseases, the molecular evolution of the pathogen is sufficiently rapid that genetic differences can accumulate between related infections on a similar time scale as disease transmission. Consequently, it can be effective to cluster cases in a high-dimensional *genetic space* in addition to clustering in physical space and time. In these studies, a case of infection is represented by a pathogen-derived molecular sequence that maps to some point in genetic space, and it may be associated with subject-level metadata such as the diagnosis date or treatment history. Clustering infections by their evolutionary relatedness is a popular method to identify and characterize subgroups with potentially elevated transmission rates [5]. For example, pairs of sequences can be clustered if the number of genetic differences between them falls below some threshold. The resulting clusters are often visualized as a network or undirected graph, where each node (vertex) represents an individual case of infection, and each edge connecting vertices indicate that the sequences of the corresponding cases are within a threshold genetic distance of each other. Sampling a group of cases that are nearly genetically identical implies that they are related through an unknown number of recent and rapid transmission events. A substantial number of genetic clustering studies have focused on the molecular epidemiology of HIV-1 [5–8]. Under current global treatment and prevention guidelines [9], greater proportions of HIV cases are being diagnosed, and new diagnoses are more frequently screened for drug resistance by genetic sequencing prior to initiating antiretroviral treatment. As a result, public health organizations are beginning to use genetic clustering methods in ‘near real-time’ to identify ongoing HIV-1 outbreaks [10, 11], to reconstruct the risk factors and etiology [12, 13], and to prioritize groups for prevention initiatives such as access to pre-exposure prophylaxis (PrEP) [14].

A significant and often overlooked challenge in the use of genetic clustering to identify potential outbreaks is that these methods usually require the specification of one or more clustering criteria [5, 15]. For instance, many HIV-1 studies that employ pairwise genetic clustering conventionally use an arbitrary threshold of 1.5% expected nucleotide substitutions per site [7, 16–18], a measure that adjusts for the transition bias and the possibility of multiple substitutions per site. In contrast, the United States Centers for Disease Control and Prevention (US-CDC) currently man-dates a stricter pairwise distance threshold of 0.5% [19]. In some cases the selected threshold is informed by the expected divergence between HIV-1 sequences sampled longitudinally from the same patient [13, 20] — however, this empirical distribution can vary substantially among subjects [21] and may be influenced by the extent of clinical follow-up. Population studies from other regions such as Botswana [22] and South Africa [23] have used substantially higher distance thresh-olds (*≥*4.5%) that vary among HIV-1 subtypes [24, 25]. Furthermore, simulation-based studies [5, 22, 26] have demonstrated that clustering is highly sensitive to the sampled proportion of the infected population. Given the known differences in the empirical distributions of HIV-1 genetic distances among populations, as well as the significant global disparities in prevalence and access to testing and treatment, it is urgently necessary to establish an objective, quantitative framework for optimizing a clustering method to the target population.

Here we propose that the most useful approach to select clustering criteria is to base this decision on our ability to predict where the next cases will occur. A high, permissive clustering threshold tends to result in a single cluster that comprises the majority of known cases. The next cases are proportionately more likely to connect to this cluster simply because it is large, but its size will also average out the individual- and group-level attributes that are informative for predicting the next cases. Put another way, a single large cluster is not likely to confer a public health benefit because it is akin to prioritizing the entire population. Conversely, setting a low, strict clustering threshold results in a large number of small clusters. This increases the variation of attributes among clusters, resolving greater information. As cluster sizes continue to decline with progressively lower thresholds, however, the variation in attributes among clusters is less associated with the emergence of new cases — in other words, the distribution of new cases among clusters becomes increasingly random. This trade-off is analogous to the modifiable areal unit problem (MAUP), a concept in spatial statistics first fully conceptualized by Openshaw and Taylor in 1979 [27]. Areal units are derived from a partition of a geographic range by drawing boundaries that separate households or neighbourhoods. The MAUP formally recognizes the inconsistency of statistical associations with changing boundaries. For example, aggregating units into larger spatial units, such as cities or countries, can prevent an investigator from detecting a strong association between water quality and gastrointestinal illness [28].

To address the MAUP in the context of genetic clustering and public health, we develop an information criterion-based framework inspired by work from Nakaya [29]. The objective of our framework is to identify the clustering criteria that maximizes the information content of the resulting clusters for forecasting where the next cases will occur. We evaluate our approach on anonymized HIV-1 sequence data from three populations, and demonstrate how this framework can also be used to select between predictive models of cluster growth that utilize different cluster attributes. Furthermore, we examine the problems associated with the arbitrary selection of clustering criteria or applying criteria from one population to another, and evaluate the stability of information-optimzed criteria for a given population over time.

## Methods

### Data collection and processing

From the public GenBank database (https://www.ncbi.nlm.nih.gov/genbank), we obtained *n* = 809 anonymized HIV-1 *pol* sequences that were sampled in northern Alberta, Canada, between 2007 and 2013 [30]; and *n* = 1, 653 sequences collected in Seattle, USA, between 2000 and 2013 [31]. We also obtained *n* = 2, 779 anonymized HIV-1 *pol* sequences from the Vanderbilt Comprehensive Care Clinic in Nashville, which were sampled from the Middle Tennessee region between 2001 and 2015; these sequences were annotated with meta-data including both years of sample collection and diagnosis [32]. Each data set was manually screened to remove all sequences corresponding to HIV-1 subtypes other than subtype B, and to remove repeated samples from the same individual. We then filtered each data set to remove any sequences with a nucleotide ambiguity above 5%, which affected 1 sequence from each of the northern Alberta and Seattle data sets, and 163 sequences from Tennessee. Given the relatively small number of sequences collected in part of 2013 for the Seattle dataset (*n* = 35, Figure 1), we excluded this year to maintain a consistent sampling rate. We retrieved the sample collection dates for Seattle and North Alberta by querying GenBank with the respective accession numbers and extracting this information from the XML stream returned from the server using the BioPython module [33] in Python. Next, we used an open-source implementation of the Tamura-Nei [34] genetic distance in C++ (TN93 version 1.0.6, https://github.com/veg/tn93) for each data-set to compute these distances between all pairs of sequences. All other options for the TN93 analyses were set to the default values.

**Figure 1:**
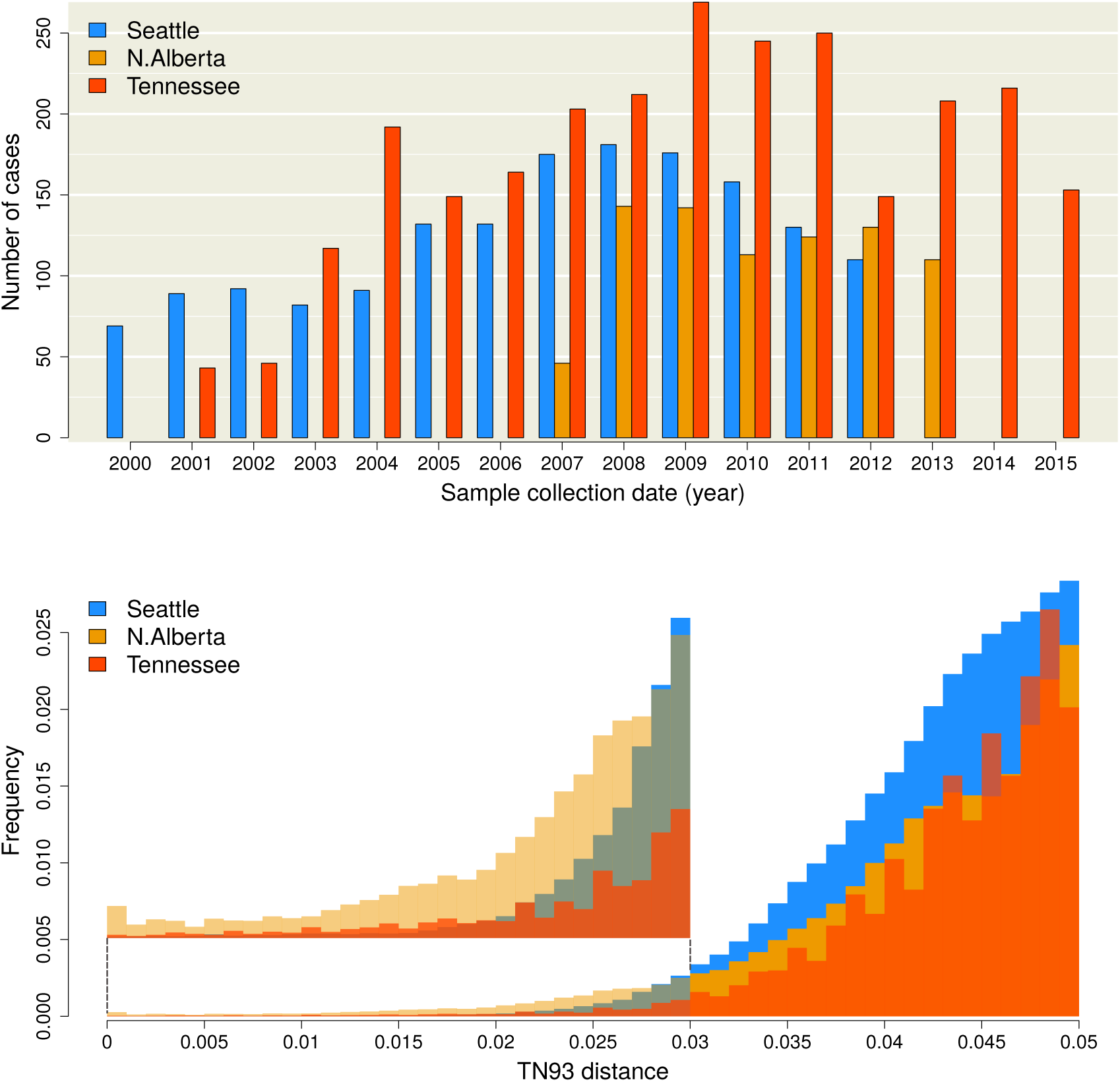
(top) Distribution of sample collection years for the Seattle (blue), northern Alberta (orange) and Middle Tennessee (red) data sets. Absent bars indicate that no sampling was carried out in the respective years, and does not reflect an absence of cases. (bottom) Histograms of Tamura-Nei (TN93) genetic distances among pairwise comparisons of HIV-1 sequences. The height of each bin has been rescaled to reflect the total number of pairwise comparisons, for which the majority were censored from the data. An expanded section of the barplots in the range (0, 0.03) is provided as a figure inset to clarify differences among the distributions.

### Defining clusters

Using a custom script in the R programming language, we generated an undirected graph 𝒢 = (𝒱, 𝓔) from the TN93 output of each data set, where the set of vertices 𝒱 represents individual cases (assuming one sequence per case) connected by edges in the set 𝓔. Every edge between vertices *v* and *u*, denoted *e*(*v, u*) ∈ 𝓔, was weighted with the TN93 distance between the respective sequences, which we denote by the edge attribute *d*(*v, u*). In addition, each vertex *v* ∈ 𝒱 carries a temporal attribute *t*(*v*), which may represent the year of diagnosis or sample collection. Note that we are not limited to analyzing dates at the level of years and can utilize more precise time intervals, *e.g.*, quarters or months, given the availability of these data. For a given clustering threshold *d*_max_, we obtained a spanning subgraph *G_d_* = (𝒱, *E_d_*) from 𝒢 that results from filtering the complete list of *n*(*n−* 1)*/*2 edge weights, such that *E_d_* = *{e*(*v, u*) ∈ 𝓔 : *d*(*v, u*) *≤ d*_max_*}*. The subset of sequences with the most recent collection time-point was specified as *V* = *{v* ∈ 𝒱 : *t*(*v*) = *t*_max_*}*, such that the total number of new cases is *|V |*. In other words, sampling time cuts 𝒱 into disjoint vertex sets *V* and *V^c^*, where *V^c^* is the complement of *V* (*V ^c^ ∪V* = 𝒱) and contains all known cases over time *t < t_max_*. Later it will be useful to refer to the subset of edges in *E_d_* that connect a vertex in *V^c^* and a vertex in *V*, which we denote as *E* = *{e*(*v, u*) : *e ∈ E_d_, v ∈ V^c^, u ∈ V}*. The set of edges in *E* can also be interpreted as edges in a bipartite subgraph comprising parts *V^c^* and *V*.

A clustering method defines a partition on the known cases *V^c^* into a set of clusters *{C*_1_*,C*_2_*, …, C_n_}* such that *C_i_ ∩ C_j_* = ø for all *i* ≠ *j* and 1 *≤ i, j ≤ n*; and such that the union of all clusters recovers the entire set: 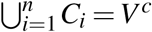. Note that this definition does not strictly require the existence of edges, which we use to represent genetic similarity, but can be adapted to any method that defines a partition on the database of known cases. For our analysis, clusters were defined as the connected components of *V^c^*, meaning any pair of vertices within the same cluster (*v, u ∈ C_i_*) are connected by at least one path (a sequence of edges), any pair of vertices in different clusters are not connected by any path, and single cases can count as their own cluster of size one.

### Modeling growth

We define total cluster growth *R* as the number of new cases in *V* adjacent (connected by an edge) to any known case in *V^c^*, where *R ≤ |V |*. To resolve the event that a new vertex in *V* is adjacent to vertices in more than one cluster, we reduced the subset of edges between *V* and *V^c^*, maintaining only the edges with minimum weight per vertex in *V*. If more than one edge to a given vertex *u ∈ V* had exactly the same minimum weight, then we selected one edge at random.

We formulated two predictive models to generate estimates of growth for the *i*-th cluster *C_i_*, which we denote respectively as the null model and the weighted model. The null model requires less information by postulating that each cluster is expected to grow in proportion to its current size, prior to the addition of new cases, as a fraction of the entire population of known cases. For example, a cluster that comprises half of all known cases is predicted to accumulate half of new cases that are adjacent to any cluster. Expressed as a Poisson regression model, the expected growth of *C_i_* given total cluster growth *R* is given by:

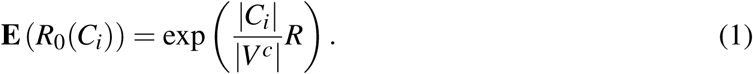

where we use boldface **E** to denote the expectation (and distinguish it from our edge set notation *E*), and *R*_0_ is given the subscript 0 to indicate it is the predicted growth under the null model. Thus the null model does not use any individual-level attributes to predict cluster growth — it is a naïve model that assumes that the allocation of cluster-adjacent new cases in *R* is not influenced by any characteristics of those clusters other than the ‘space’ they occupy. In contrast, the weighted model assigns individual-level weights *w*(*v*) to every vertex in *v ∈ C_i_*. Expressed as a Poisson regression model, the expected growth of cluster *C_i_* under the weighted model is written:

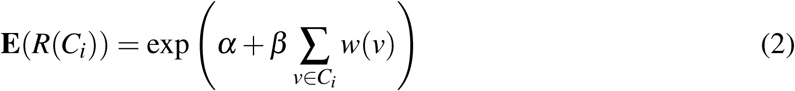

where *α* and *β* are parameters to be estimated by regression. Note that equation (2) reduces to (1) when *w*(*v*) = 1 for all *v ∈ C_i_*, *α* = 0 and *β* = *R/|V^c^|*.

For our demonstration, we weighted individual cases by their recency of sample collection or diagnosis, measured as Δ*t* = *t_max_ − t*(*v*). The predictive weight *w* of a known case of a given age Δ*t* relative to *t_max_* was based on the expected rate of adjacency (edge density) between sets of known cases separated by the same time lag. Thus, we needed to calculate the edge densities for all bipartite graphs 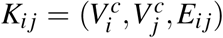 where *V^c^_i_* = *{v ∈ V^c^* : *t*(*v*) = *i}* and *j − i* = Δ*t*. For compatibility with our definition of cluster growth, we removed bipartite edges from *E_i j_* so that the maximum degree size for any vertex *v ∈ V^c^_j_* was 1, where the remaining edge minimized the edge weight *w*(*u, v*) for all *u* given *v*. We use 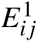 to denote this reduced set of bipartite edges. This had the effect of reducing the maximum possible number of bipartite edges in *K_ij_* from *|V_i_||V_j_|* to *|V_j_|*. We refer to the set of all bipartite graphs for a given time lag as *K*(Δ*t*) = *{K_i j_* : *j > i, j − i* = Δ*t}*. Thus, the expected edge density *ρ* given Δ*t* is:

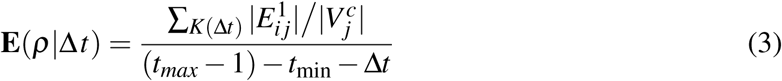

For this model, we assume that the edges in 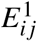 are independent and identically distributed binary outcomes. Furthermore, we expect the probability of this outcome to decay with increasing Δ*t*.

Hence we used logistic regression to estimate *ρ* as a function of Δ*t*:

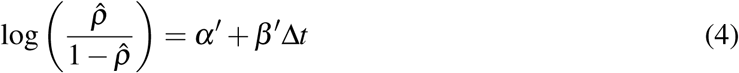

where *α*′ and *β*′ are parameters to be estimated by regression. The simplest use of this information would be to set the weight of a known case to its predicted edge density given its time lag Δ*t*; *i.e.*, 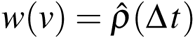.

The weighted model can be extended to employ a linear combination of additional individual-level attributes (*e.g.*, plasma viral load) and/or graph attributes that are parameterized from bipartite subgraphs on *V^c^*. For example, we added an additional measure, deg(*v*), that represents the average degree of known cases from the same time point as *v*. A high mean degree in the graph for a given threshold *w*_max_ can reveal specific time points with an unusually dense concentration of cases in genetic space, which may be caused by a period of increased sampling effort or a past outbreak. Without making some adjustment, this period can have a disproportionate influence on the association of edge density on Δ*t* as estimated by equation (4). Thus we also evaluated a weighted model substituting the following weighting formula into equation (2):

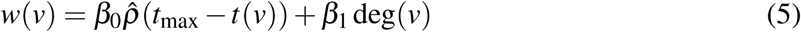

where the coefficient *β* has been brought into the summation over individual known cases. In this model, deg(*v*) is a graph-level attribute that controls for the confounding effect of variation in degree size among years of diagnosis.

### Evaluating cluster thresholds

For each data set, we segregated all HIV-1 sequences that were sampled in the most recent year as new cases comprising the set *V*. Next, we extracted the observed cluster growth outcomes *R*(*C_i_*) and individual case weights *w*(*v*) at 51 different cluster-defining distance thresholds, ranging from *d*_max_ = 0 to *d*_max_ = 0.04 in steps of 8 *×* 10*^−^*^4^. We fit the null and weighted models described by Equations (1) and (2) to the resulting distributions of cluster growth. The Akaike information criterion (AIC), which penalizes likelihood for the number of model parameters, was recorded for each regression model [35]. The ‘generalized AIC’ (GAIC) is simply the difference in AIC between models, and has been proposed as a key quantity for resolving the modifiable areal unit problem [29]. Cutoffs with a negative GAIC indicate that the weighted model explains the data more effectively than the null model, and the magnitude of GAIC quantifies that difference in effectiveness. We define the optimal distance threshold as the value *d*_max_ associated with the lowest (most negative) GAIC. The GAIC was evaluated for the weighted models using either dates of sample collection or diagnosis to compute Δ*t* for the Middle Tennessee data set for which both dates were available. Since some dates were missing data, we normalized the number of new cases to *|V |* = 125 for both analyses to ensure that growth rates were comparable. After this step, the total numbers of cases was reduced to *n* = 2, 015 and *n* = 2, 588 for the diagnostic and sample collection analyses, respectively. We note that the resulting weighted models are not being compared directly; instead, they are compared to the null models for their respective data sets.

We repeated the cluster threshold evaluation on progressively censored subsets of the Seattle and Tennessee data to evaluate the consistency of the GAIC-optimized thresholds over time. This was accomplished by removing cases from the most recent year to a maximum of four years, and obtaining the GAIC measurements for all values of *d*_max_ for the remaining data at each step. Because of the limited size and temporal range of the northern Alberta data set, we did not use it for this sensitivity analysis. We also re-ran the experiment on random subsets of cases from the complete Seattle and Tennessee data sets, creating a total of 90 subsets; 30 sub-sets at 3 different proportions of the original total, 80%, 60%, and 40%. For each resampling proportion, we obtained the kernel density for the optimal cutoff location over the 30 replicate samples and the GAIC measurements obtained by a smoothed function from all 90 resampled GAIC runs.

## Results

### Study populations

A total of *n* = 5, 010 HIV-1 sequences and sample collection dates were obtained from published studies in Seattle (*n* = 1, 591) [31], northern Alberta (*n* = 803) [30], and Middle Tennessee (*n* = 2, 616) [32] respectively. In addition, dates of HIV-1 diagnosis were available for a total of 2, 527 cases in the Tennessee data set. The distributions of sample collection dates are summarized in Figure 1 and the direct comparison between diagnostic and collection year distributions for the Tennessee data set can be found in supplementary Figure S1. The lowest mean sampling rate was obtained in northern Alberta, with 114.7 cases per year, compared to 122.4 and 174.4 cases/yr for Seattle and Tennessee respectively. Cases from the final years of sampling were omitted to adjust for studies being terminated before the end of the calendar year. For instance, there were only 35 cases sampled in 2013 in the Seattle data set, where sample collection was ended on March 2013 [31] — since the cases in the final year were reserved to evaluate the predictive models, an artificially low sample size would have a disproportionate influence on model validation. Hence, we proceeded with 110 cases collected in 2012, 110 cases from 2013, and 153 cases from 2015 for Seattle, northern Alberta and Tennessee, respectively. We refer to the cases sampled in these final years as ‘new cases’, and those sampled in the preceding years as ‘known cases’.

For each data set, we calculated the Tamura-Nei (TN93 [34]) genetic distance for every pair of sequences. This distance, which adjusts for differences in the mean rates among nucleotide transversions and the two types of transitions, is the basis for clustering in the HIV-TRACE program [36] that is employed by the U.S. Centers for Disease Control and Prevention for public health surveillance [37]. Although the northern Alberta data set comprised a smaller number of sequences, the lower tail of its TN93 distribution contained relatively higher numbers of pair-wise distances than the other two data sets (Figure 1). For instance, the TN93 distance at the 1% quantile was 0.013 in northern Alberta, while the same quantile was roughly twice this distance for Seattle (0.026) and Tennessee (0.023). Overall, these distributions were significantly different (Kruskal-Wallis test, *P <* 10*^−^*^15^).

### Adjacency of cases decays with time lag

We generated a sequence of graphs at varying TN93 distance thresholds for each data set, where each vertex represents a known case (sampled or diagnosed prior to the final year) and an edge indicates that the corresponding pairwise distance is below the threshold — in graph theory, the cases are said to be ‘adjacent’. Thus, each distance threshold defines a different partition of known cases into clusters, where a cluster may consist of only a single known case. Our objective is to determine which threshold results in the most information-rich partition of known cases for predicting where new cases will arise. As we will demonstrate below, there is no information value in either extreme of a single giant cluster or the complete atomization of cases into singular clusters. To quantify the information loss associated with different partitions, we compared two predictive models. First, we fit a null model that assumes the probability that a new case appears in a cluster (*i.e.*, cluster growth) is only influenced by the number of known cases in the cluster, *i.e.*, the cluster size. This is equivalent to assuming that every known case is equally likely to be adjacent (connected by an edge) to the new case. Second, we fit a weighted model where the probability of cluster growth is predicted by some linear combination of individual-level attributes among the known cases in the cluster.

For example, we hypothesize that the probability that a new case is adjacent to a known case declines with an increasing time lag between their respective sample collection dates. To investigate this effect, we plotted the observed densities of edges at a threshold *d*_max_ = 0.04 between sets of known cases sampled in different years (Figure 2). These plots confirm that edge densities decline significantly with increasing time lag (Δ*t*), which we measured by fitting binomial regression models (equation 4). Specifically, the estimated effect of Δ*t* on the log-odds of a bipartite edge was *−*0.42 (95% C.I. = *−*0.45*, −*0.39) year*^−^*^1^ for the Seattle data and *−*0.40 (*−*0.48*, −*0.32) year*^−^*^1^ for northern Alberta. The coefficient of determination for the respective models was *R*^2^ = 0.70 and 0.58. For the Tennessee data set, the effect of time lag was lower than the other data sets (*−*0.19 (*−*0.21*, −*0.18) year*^−^*^1^; *R*^2^ = 0.41). In general, lowering *d*_max_ reduced the observed bipartite edge densities as fewer edge weights passed the threshold. Nevertheless, the negative associations between Δ*t* and the log-odds of bipartite edges were robust to varying *d*_max_ (Supplementary Figure S2). These results supported the use of ‘case recency’ (the time lag between sample collection dates) as an individual-level predictor of a new case joining a cluster of known cases.

**Figure 2:**
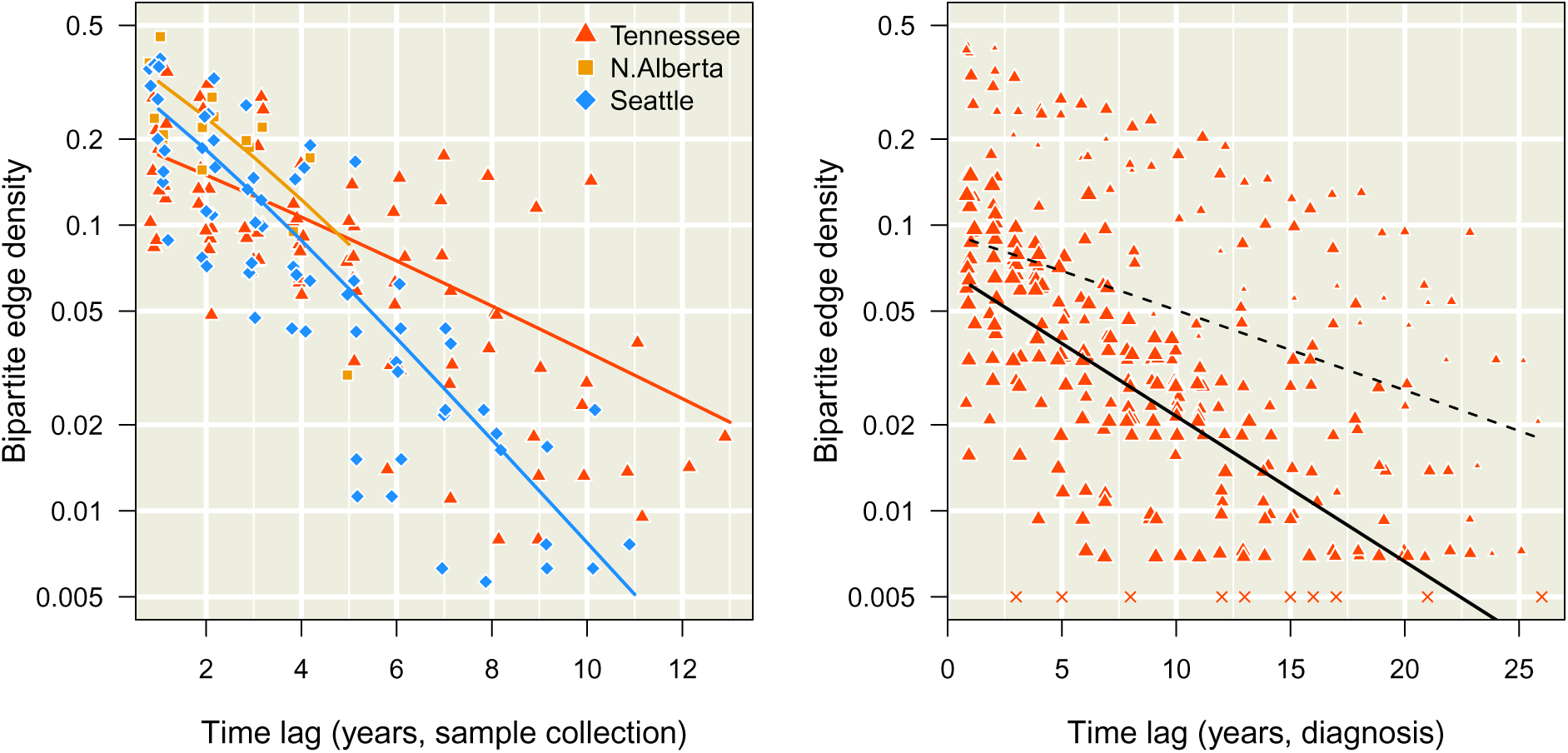
Decay in bipartite edge densities with increasing time lags. Each point represents a bipartite graph at a given time lag (difference in years, *x*-axis), coloured with respect to the data set (see inset legend). We added random noise along the horizontal axis to separate overlapping points. The log-transformed *y*-axis represents the frequency of edges between cases in different time points below the threshold *d_max_* = 0.04 (Equation 3). **(left)** Decay of edge densities with respect to dates of sample collection. Each trend line summarizes the binomial regression model (Equation 4) for each data set. **(right)** Decay of edge densities with respect to dates of HIV diagnosis in Middle Tennessee. Each point was scaled in proportion to the mean degree size among known cases in the earlier time point. Bipartite graphs without any edges are represented by × marks at the arbitrary value of 0.005, since zero cannot be displayed on a log-transformed axis. The dashed line indicates the fit of the binomial regression to these data, while the solid line indicates the fit of an extended model (Equation 5) that controls for variation in the mean degree size.

Since dates of HIV-1 diagnosis were also available for the Tennessee data set, we applied the same binomial regression to known cases separated by their years of HIV diagnosis rather than by sample collection (Figure 2, right). We noted that the decline in adjacency rates with time lag was distorted by unusually high edge densities involving cases diagnosed in 1982, 1984 and 1986, which resulted in a much smaller but significant effect of Δ*t* (*−*0.067 (95% C.I. = *−*0.074*, −*0.060) year*^−^*^1^; *R*^2^ = 0.016). This motivated the use of an extended binomial regression model (Equation 5) to control for variation in degree size among years of diagnosis for known cases. We found that controlling for variation in degree sizes among years substantially improved the fit of the regression model to the time lags in diagnosis dates (*R*^2^ = 0.37, Δ*AIC* = *−*351.4). Moreover, this extended model conferred significantly improved fits to sample collection dates for all three data sets (Δ*AIC* = *−*7.5, *−*12.8 and *−*171.6 for Seattle, northern Alberta and Middle Tennessee, respectively). Therefore, we used the extended binomial regression model for our subsequent analyses to predict the distribution of new cases among clusters.

### Trade-off between case coverage and cluster information

Figure 3 illustrates the effect of relaxing the threshold *d*_max_ on the number of new cases that are adjacent to one or more known cases, which we denote by *R*. A new case that is not adjacent to any known case cannot be anticipated by any clustering method. When *R* approaches the total number of new cases (denoted as *|V |*, where *V* is the set of vertices in the final year) we say that the clusters have a high *case coverage*. As expected, decreasing *d*_max_ reduced *R* as a progressively greater number of edges were excluded, which would limit the utility of clustering for public health surveillance. The accumulation of *R* with increasing *d*_max_ was visibly slower for Seattle than the other data sets (Figure 3, left), which was not anticipated by our comparisons of the overall distributions of TN93 distances among data sets (Figure 1, bottom). In addition, Figure 3 (left) summarizes how the number of clusters with at least one edge to a new case (the number of active clusters) initially tracks the accumulation in *R* with increasing *d*_max_. Thus *R* sets an upper limit to the number of active clusters; these numbers can only be equal if each new case is uniquely adjacent to its own cluster, as we observed for the Seattle data set for cutoffs below *w*_max_ = 0.0072 or the Tennessee data at cutoffs below *w*_max_ = 0.008. As *d*_max_ continues to increase, the number of active clusters peaks and begins to decline towards 1. This outcome reflects the gradual accretion of cases into a single giant cluster. Substituting year of diagnosis for sample collection dates in the Tennessee data resulted in a slight reduction in *R* and a modest increase in the number of active clusters, which we attribute to a more uniform distribution of new cases across clusters.

**Figure 3:**
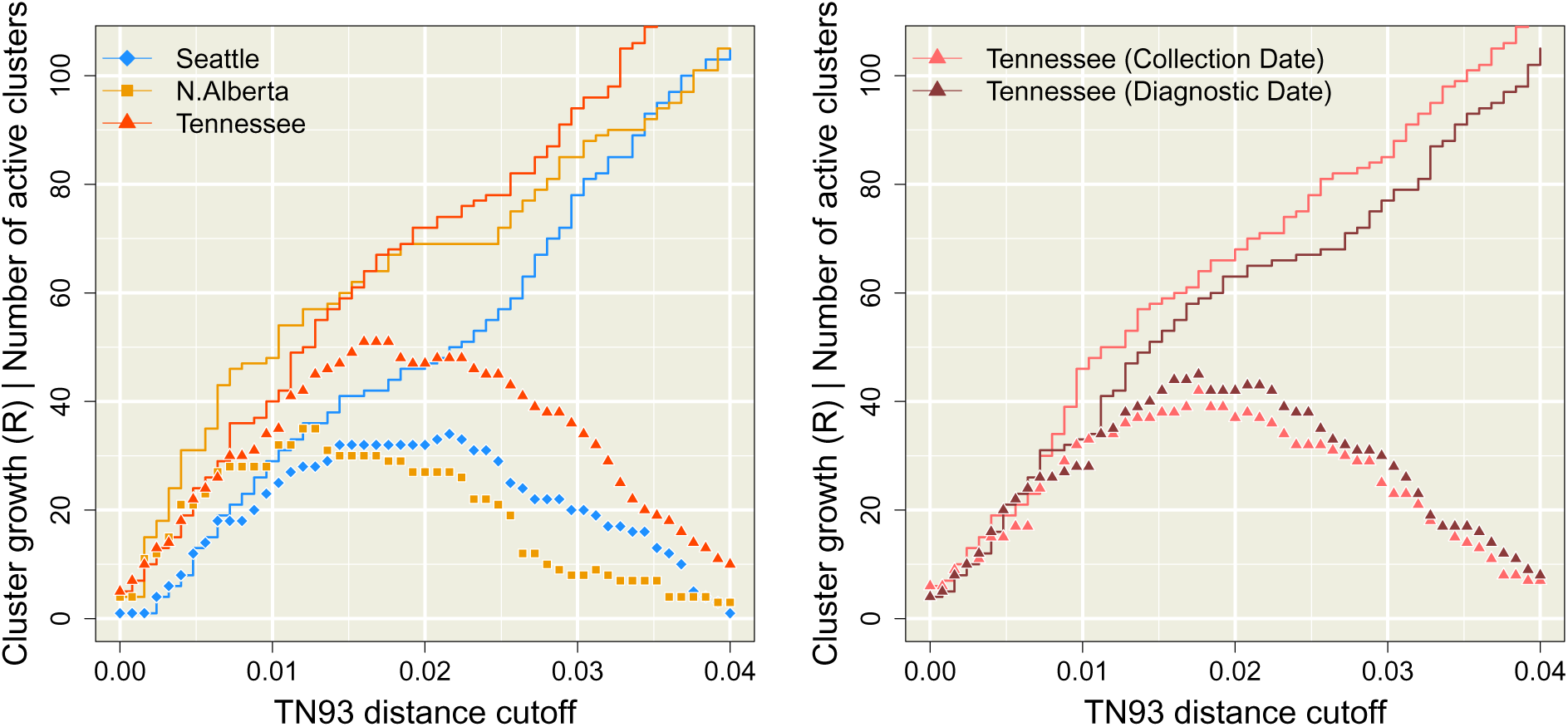
Distribution of new cases among clusters as a function of the Tamura-Nei (TN93) distance clustering threshold (*d*_max_, *x*-axis). The solid lines represent the total number (*R*) of new cases adjacent to clusters of known cases. The points correspond to the number of clusters of known cases with edges to new cases, which we refer to as ‘active’ clusters. **(left)** This plot summarizes the trends obtained when cases are stratified by year of sample collection. Trends in *R* and numbers of active clusters are coloured with respect to data set (see inset legend). **(right)** This plot contrasts the trends obtained from the Tennessee data set when cases were stratified by year of sample collection (lighter red) versus the year of diagnosis (darker red). Note that the collection date trend is not identical to the trend in the left plot because we downsampled cases to match the availability of diagnosis dates.

### Obtaining GAIC

The results in the preceding section imply that there exists an intermediate value of *d*_max_ that optimizes the trade-off between case coverage and the number of active clusters, where both quantities have a significant impact on the information content of clusters for public health. We propose that the best criterion for optimizing a clustering method is our ability to predict where the next cases will occur among the resulting clusters. Specifically, we adapted the generalized Akaike information criterion (GAIC [29]) to select the optimal threshold. Our implementation of the GAIC is a comparison between two Poisson regression models, where the count outcomes are the numbers of new cases adjacent to each cluster. In the null model, we assume that the rate parameter is proportional to cluster size as a fraction of all known cases (Equation 1), assuming no variation among individual known cases. In the weighted model, the rate parameter is the total weight of known cases in the *i*-th cluster, where each weight can be calculated from a linear combination of individual- or group-level attributes. For our analysis, we weighted cases by their predicted edge densities from the extended binomial model (equation 5).

Figure 4 (left) summarizes the distributions of GAIC for varying *d*_max_ for each data set. We observed that GAIC tended to be near zero for relaxed thresholds (*d*_max_ *≥* 0.03), which indicated that the ability of the weighted model to forecast new cases was indistinguishable from the null model. At these high thresholds, the majority of known cases tended to become grouped into a single large cluster, thereby homogenizing any individual-level variation that could be used by the weighted model to predict the distribution of new cases. The minimum (most negative) GAIC was obtained *d*_max_ = 0.0104 for northern Alberta and *d*_max_ = 0.0160 for both Seattle and Tennessee. These minima identified the optimal thresholds that maximized the difference in information loss between the weighted and null models. As we continued to decrease *d*_max_ past these optima, the GAIC trends returned to the zero line and even increase sporadically into positive values where the weighted model was *worse* than the null model. At these low values of *d*_max_, the disintegration of clusters into large numbers of singletons disrupts the covariation between individual attributes and the distribution of new cases. Furthermore, lowering *d*_max_ also leads to a reduction of case coverage as shown in Figure 3. At the respective optimal thresholds, less than half of new cases were adjacent to clusters of known cases (38.2% for Seattle, 49.1% for Northern Alberta and 41.8% in Tennessee).

**Figure 4:**
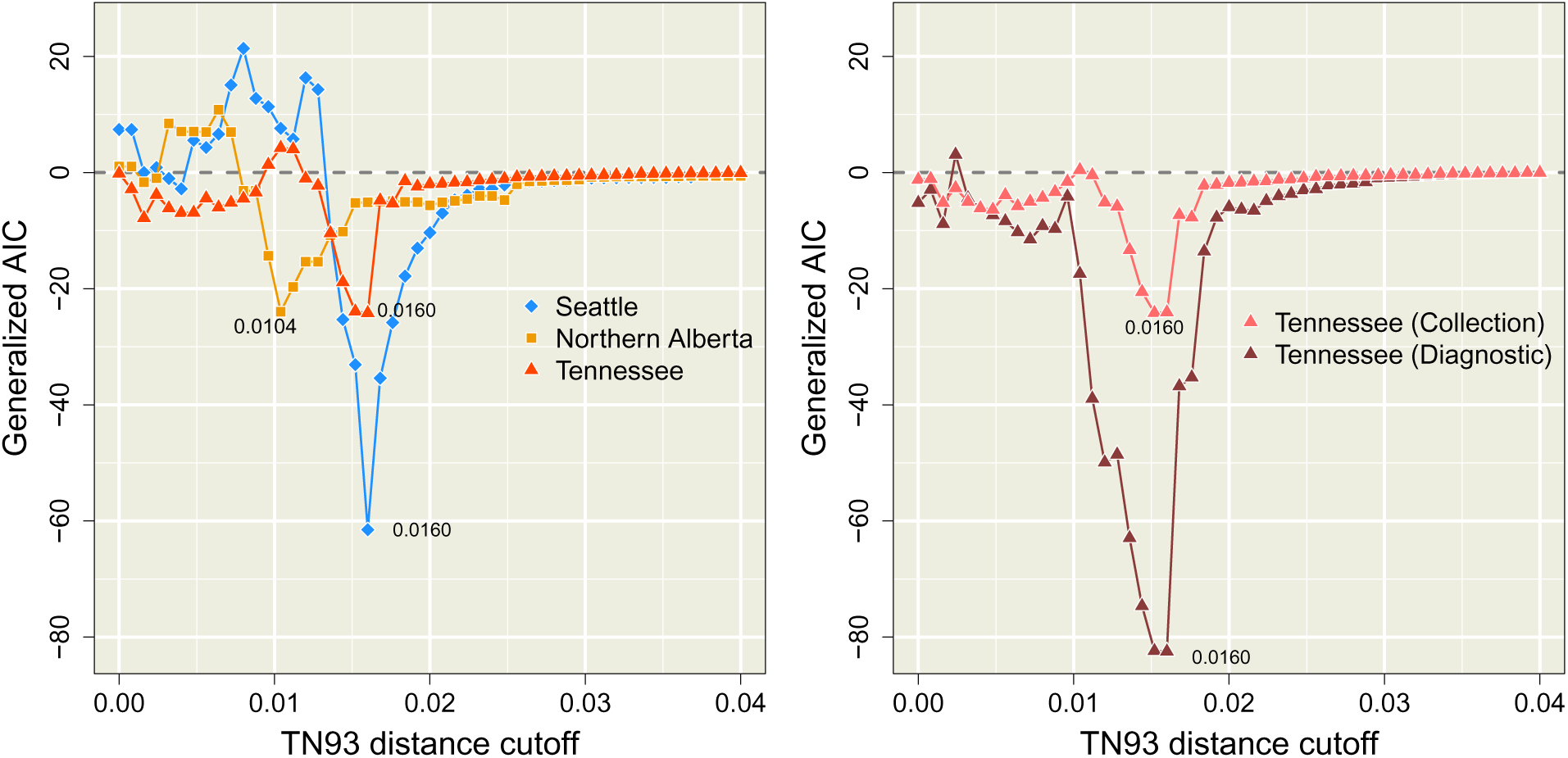
The GAIC relative to cutoff *w_max_*. This is calculated as the difference in AIC between 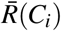 fit to *R*(*C_i_*) minus 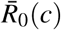 fit to *R*(*c*). Negative values from a given *w_max_* imply that the growth of that particular set of clusters is better predicted by 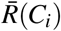 than 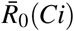. **(left)** The three complete data sets using collection dates. **(right)** The Tennessee data sets filtered for direct comparison between diagnostic and collection date.

The GAIC also provides a framework for variable selection. For instance, the GAIC obtained for the Tennessee data when cases were stratified by year of diagnosis was substantially lower than the values obtained with sample collection dates over wide range of *d*_max_ (Figure 4, right). The optimal threshold identified by the minimum GAIC coincided for both sets of dates (*d*_max_); how-ever, the GAIC for diagnosis dates was substantially more negative (ΔGAIC = *−*58.3), indicating a more effective use of cluster information. For larger values of *d*_max_ *>* 0.02, the stratification of cases by either set of dates was irrelevant as they collapsed into a single giant cluster, such that both GAIC trends converged to zero. For smaller values of *d*_max_ *<* 0.01, both trends became more erratic with the reduced adjacency of new cases and increasingly stochastic composition of smaller clusters.

The graphs for these optimal cutoffs are summarized in Figure 5. In the Seattle graph, the largest cluster (Se1) comprising 34 known cases was adjacent to 2 new cases. The sample collection years associated with this cluster range from 2000 to 2012 with a mean of 2006.8 (interquartile range, IQR = 2006 – 2009). In contrast, the second largest cluster (Se2) comprised only 10 known cases that were sampled more recently with a mean of 2009.6 (2008 – 2011), and accumulated 6 new cases. Similarly, the largest cluster in the northern Alberta graph (NA1) comprised 22 cases of which none were new, with a mean sample collection year of 2009.2 years (2008 – 2011). This contrasts a smaller cluster of 10 known cases (NA2), of which 5 were collected in 2012 (mean 2010.5, IQR = 2009 – 2012), that gained 12 new cases in 2013. Finally, we observed a large cluster (Tn1) of 72 known cases in the Tennessee data set with only one new case and a mean sample collection year of 2007.6 (IQR 2005 to 2010). In contrast the cluster labelled Tn2 comprised 39 known cases with a mean collection year of 2012 (IQR 2010, 2013) and 3 new cases. These simple examples illustrate the effect of optimizing the clustering threshold on the covariation of new case counts and the recency of known cases among clusters. As predicted by our model analysis, the effect of variation in case recency among clusters on the distribution of new cases is even clearer for the graph depicting clusters of cases stratified by year of diagnosis rather than sample collection (Supplementary Figure S3).

**Figure 5:**
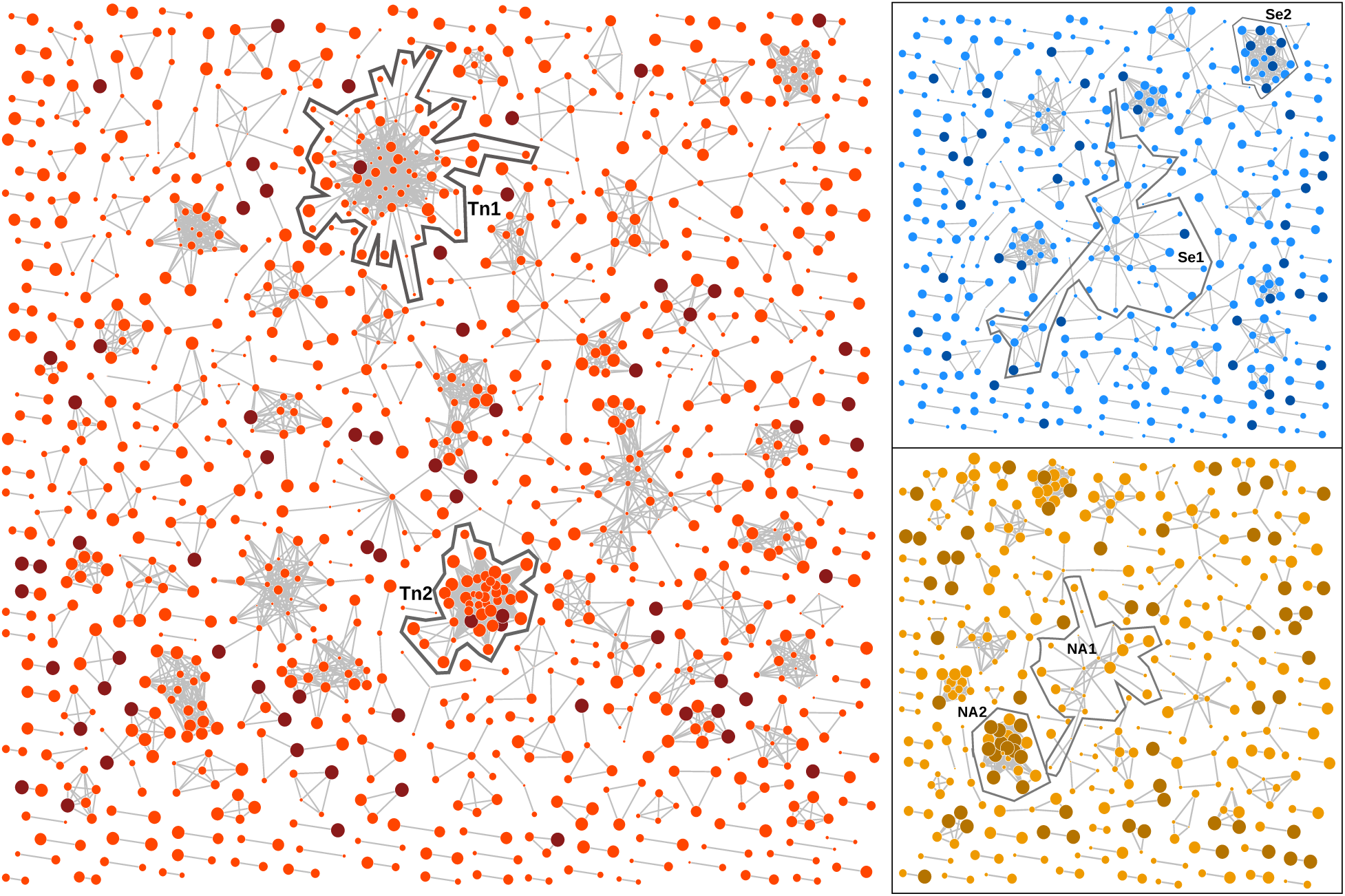
Network visualisations of the graphs from the Middle Tennessee data (red), Seattle (blue) and northern Alberta (orange) data sets obtained at the respective GAIC-optimized distance thresholds, rendered using an implementation of the Kamada-Kawai [38] algorithm. Vertices that corresponded to new cases were coloured in a darker shade, and the width of each vertex was scaled to the sample collection year with more recent cases drawn at a larger size. The vertices with zero degrees (singletons) were not drawn for clarity. Fast growing clusters and/or large clusters have been highlighted and labelled.

### Robustness of GAIC optimization

The difference in optimal clustering thresholds between northern Alberta and the other sites implies that a globally optimal threshold does not exist. However, variation in thresholds may also be a stochastic outcome due to incomplete sampling. To measure the effect of sampling variation, we repeated our GAIC analysis on random subsamples of the Seattle and Tennessee data sets to 40%, 60% and 80% of the data (30 replicates each). The comparably low number of cases per year in the northern Alberta data set precluded subsampling. The results for the Seattle data set are summarized in Supplementary Figure S4. As expected, we observed shallower GAIC trends and more variable GAIC-optimized thresholds with decreasing sample size. However, the optimal thresholds remained clustered around the original value *d*_max_ = 0.016, with only 4 (13%) replicates with optima below *d*_max_ *<* 0.015 at 40% subsampling (*n* = 636, comparable to the northern Alberta sample size of *n* = 803). We obtained similar results on random sub-samples of the Tennessee data set, except that the optima were more robust for the data stratified by diagnosis dates.

Finally, to assess the stability of GAIC-optimized thresholds over time, we generated additional subsamples by progressively right-censoring the Seattle and Tennessee data sets. If the information content of pairwise TN93 clusters is stable over time, then we should expect the optimized threshold from fitting models to ‘new cases’ in a given year should be similar to the optimized threshold with an additional year of case data. Figure 6 summarizes our results for the Tennessee data set where the four most recent years were progressively censored by dates of sample collection (left) and diagnosis (right), respectively. Results for the complete Tennessee data set (including cases with missing diagnosis dates) and the Seattle data set are provided in the Supplementary Figure S7. In general, we observed that the optimal threshold identified by the minimum GAIC was relatively stable over time. Although the exact thresholds varied slightly, the optimal threshold from the previous year tended to confer a GAIC similar to the threshold estimated from that year. In our analysis of the Seattle data set, however, the optimal distance thresholds fluctuated between two local minima around *d*_max_ = 0.005 and 0.015. For instance, in the subset where cases were limited to 2011 (censoring cases sampled in 2012) we observed three local minima in the GAIC (*d*_max_ = 0.0024, 0.0072 and 0.0144), including one that was close to the global minimum of the previous year. These results suggest that the larger database of cases sampled in Middle Tennessee confer increased robustness to sampling variation, although we cannot rule out site-specific effects.

**Figure 6:**
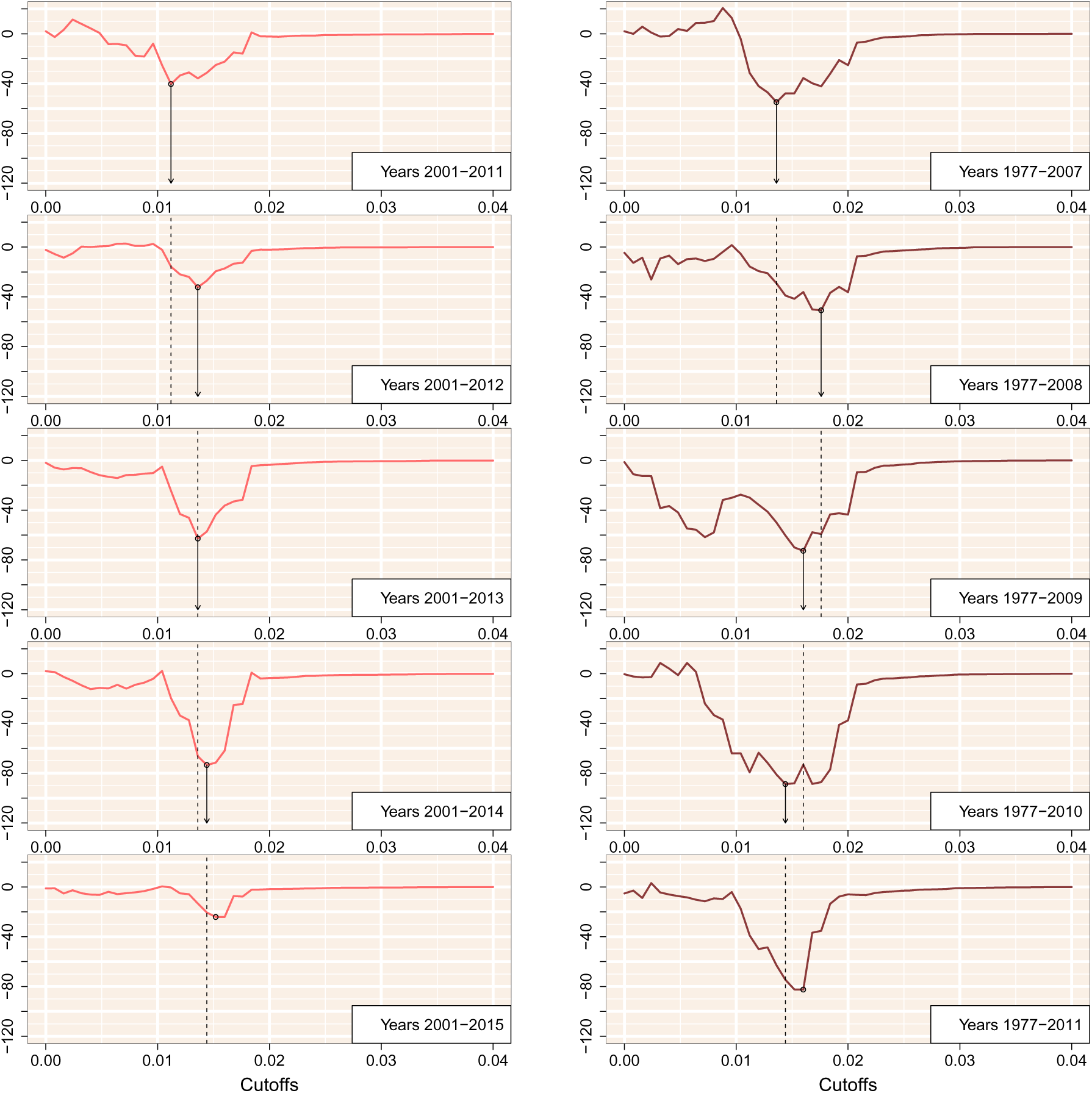
GAIC plotted against the cutoff threshold for five progressively right-censored subgraphs with respect to sample collection (**left**) and diagnosis (**right**) for the Middle Tennessee data set. The minimum GAIC for each plot is indicated by a circular mark. Arrows and dashed liness indicate where the GAIC-optimized threshold from the previous result would carry over to the subsequent year.

## Discussion

Our results demonstrate that an apparently small difference in pairwise genetic distances — for instance, between 0.5% and 1.5% — can make the difference between accurate forecasting of new cases among clusters and becoming misled by stochastic noise. Specifically, both cutoffs cited above are routinely used as customary settings in pairwise genetic distance clustering studies of the same HIV-1 subtype in the same country [18, 19, 31]. To investigate the sensitivity of clustering thresholds, we have examined three data sets of anonymized HIV-1 subtype B *pol* sequences that were collected in different regions of North America within similar time frames. All GAIC-optimized cutoffs are located in the left tails of the respective empirical distributions of pairwise distances, with no clear demarcation that might motivate the selection *a priori* of one cutoff over another (Figure 1). However, our information-based criterion reveals a stark difference between these cutoffs when we evaluate the ability of genetic clusters to ‘forecast’ the occurrence of new cases (Figure 4). This discordance is a result of a tradeoff between the coverage and predictive value of spatial information that is encapsulated by the modifiable areal unit problem (MAUP [29]). As we relax the clustering threshold, instances of cluster growth become more frequent such that aggregate effects (*viz.*, case recency and mean degree) can be distinguished against the background of stochastic effects. However, the variation in growth rates among clusters eventually becomes homogenized as they are collapsed into a single giant cluster at increasingly high thresh-olds. These two driving forces can be differentiated in their approach to an optimum, with the poor case coverage generating stochastic changes in ‘forecasting’ ability and the lower variation creating a much smoother loss of information (Figures S4, S5, and S6).

Unlike most instances of the MAUP that arise in spatial epidemiology [28, 29, 39, 40], our outcome variable (the number of new cases per genetic cluster) is directly dependent on the same parameters that reshape the partition of the spatial distribution of covarates into units. This dual dependency results in an asymptotically increasing model likelihood with increasing distance thresholds, plateauing at the point where all known cases were assigned to the same giant cluster, such that any new cases are effectively guaranteed to be adjacent to this cluster. We addressed this unique problem by formulating a null model where the predicted growth of a cluster was directly proportional to its relative size in the number of known cases. Hence, this null model provided a useful baseline that controlled for the proportionate effect of the largest cluster with increasing cutoffs, thereby enabling us to focus on the predictive value of variation in covariates among clusters.

We selected a relatively simple but established clustering method (pairwise TN93 distance clustering by components [17]) to demonstrate our new framework for evaluating clusters, based on Nakaya’s generalized Akaike information criterion (GAIC) [29]. Despite the simplicity of pair-wise clustering, it has been widely adopted in health jurisdictions around the world, including the U.S. Centers for Disease Control and Prevention [37], in part due to the growing popularity of the HIV-TRACE software package [36]. However, we hypothesize that it should be feasible to use this framework to evaluate potentially any clustering method that defines a partition on the database of known cases. For example, if we require some minimum bootstrap support value to define clusters as subtrees extracted from the total phylogenetic tree [41], then this bootstrap support threshold can represent a second dimension (in addition to a branch length threshold) to locate the minimal GAIC in combination with a distance threshold. We have also demonstrated that the GAIC can be repurposed for model selection where different linear combinations of predictor variables, such as the mean degree size in a given year of diagnosis (Figure 4), are evaluated within the Poisson regression model. Billock and colleagues [7] recently employed a similar model selection approach for pairwise TN93 clusters, although their analysis pre-specified a fixed clustering threshold of 1.5%. Thus, if sufficient metadata are available then one can use the GAIC to select more accurate predictive models of cluster growth while adjusting the clustering criteria, so the models can be evaluated at their best performance.

Collection dates in units of years are most frequently available as sample metadata in association with published HIV-1 genetic sequences. We had a strong *a priori* expectation for an association between new case adjacency and known case recency that we subsequently confirmed from these data (Figure 2). On the other hand, we recognize that samples may be collected well after the start of a new infection, due to the long asymptomatic period of HIV-1 infection and social barriers to HIV testing [42]. Although estimated dates of infection (*e.g.*, the midpoint between the last HIV seronegative and first seropositive visit dates) will inevitably be closer to the actual date of infection, the necessary information for accurate estimates are not routinely available in a public health context. We furthermore recognize that more precise dates of sample collection would likely confer greater prediction accuracy. Indeed, the granularity of time in the context of genetic cluster analysis represents an extension of MAUP that is known as the modifiable temporal unit problem [43]. While reducing the length of time intervals may produce more timely predictions, *e.g.*, new cases in the next three months instead of the next year, the accuracy of prediction will erode with progressively shorter intervals. Finally, we propose that an informative assessment on the potential value of genetic clustering for public health would be to compare the GAIC of the genetic clustering method against the value obtained from the prioritization of groups by public health experts. However, the confidential information comprising the latter case is not likely to be found in the public domain.

An important caveat to our approach is that the expected probability of a edge between specific known and new cases is very small. Consequently, our method requires a substantial number of new cases to parameterize models of the variation in edge densities among clusters and, ultimately, to discriminate between the null and weighted models. (Note that the number of cases sampled in a given year does not correspond to the annual incidence.) Given the results summarized in Figure 6, if those requirements are not met, the minimal GAIC that results from the theoretical MAUP tradeoff may be masked by the initial noise of a graph with low case coverage and the merging of clusters with increasing thresholds. The results that we obtained with the smallest data set (N. Alberta) and our sub-sampling experiments (Figures S4 – S6) imply that averaging about 50 to 100 sampled cases per year over a 6 year period should be adequate for the relatively simple models evaluated here. In addition, our comparison of sample collection versus diagnosis dates (Figures S5 and S6) implies that that the minimum sample size requirement should diminish with the addition of covariates into the weighted model that have strong associations with the distribution of new cases.

Genetic clustering is used increasingly for near real-time monitoring of clinical populations for the purpose of guiding public health activities [6, 7, 10, 37]. Our method is not specific to HIV-1, although the proliferation of clustering methods in HIV molecular epidemiology — driven by the abundance of genetic sequence data and the relatively rapid rate of evolution and low transmission rate of the virus — does make this approach particularly applicable to this virus. Similar pairwise distance clustering methods, for instance, have been used for *Mycobacterium tuberculosis* [44] and hepatitis C virus [45] to infer epidemiological characteristics from molecular sequence variation. In these cases, it may be necessary to rescale the step size/range of clustering thresholds to the expected distribution of genetic distances in order to locate the minimum GAIC. No matter what pathogen is the focus of investigation, however, it is imperative that we become more critical of our methods [5, 46]. Improperly calibrated clustering methods may otherwise prioritize false-positives, diverting limited public health resources away from subpopulations where the immediate need for prevention and treatment services was greatest.

## Acknowledgements

This study was supported by a Project Grant from the Canadian Institutes of Health Research (PJT-156178) and by an Administrative Supplement to a Center Core Grant from the Tennessee Center for AIDS Research (P30-AI110527). CC was supported by a Dr. Frederick Winnett Luney Graduate Research Award from the Department of Pathology at Western University, Canada. An early version of this work was presented at the 26th International HIV Dynamics & Evolution meeting in Cascais, Portugal. We thank John R. J. Thompson for a helpful discussion on applications of spatial statistics to model forest fire risk, which motivated us to look beyond autoregressive models; and Dr. Tomoki Nakaya for encouraging feedback at an early stage of this study.

Data available at Genbank accession numbers KY034691–KY037792, KU190031–KU190839, and MH352627–MH355541.

## Supplementary Figures

**Figure S1:**
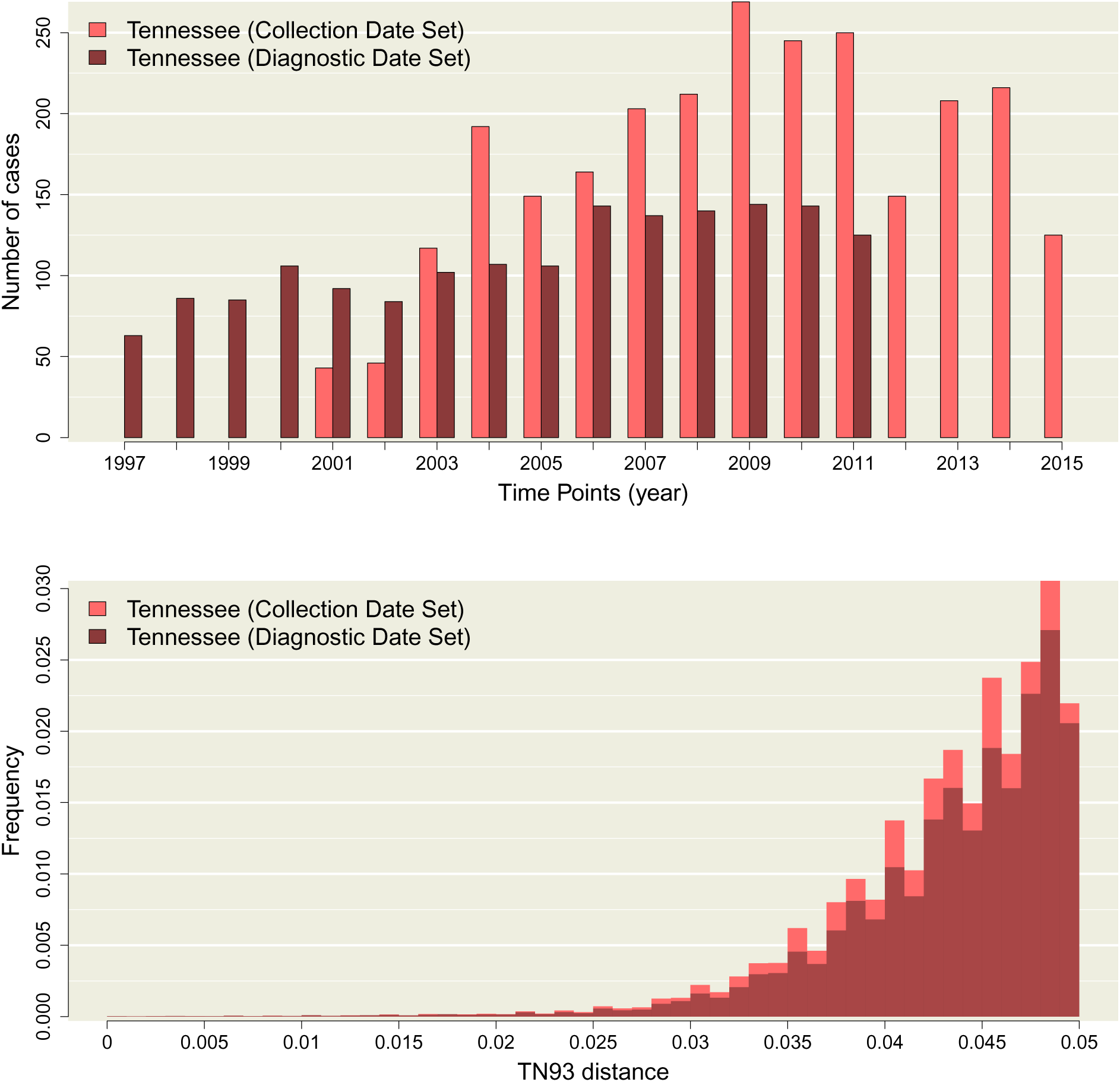
**(top)** Distribution of sample collection years for the Tennessee data set (pink) and diagnostic years for the same location (red). Absent bars indicate that no sampling was carried out in the respective years, and does not reflect an absence of cases. Each data set was filtered such that **(bottom)** Histograms of Tamura-Nei (TN93) genetic distances among pairwise comparisons of HIV-1 sequences from Tennessee (pink) with collection dates compared to diagnostic years (red) for the same location. The height of each bin has been rescaled to reflect the total number of pairwise comparisons, for which the majority were censored from the data.

**Figure S2:**
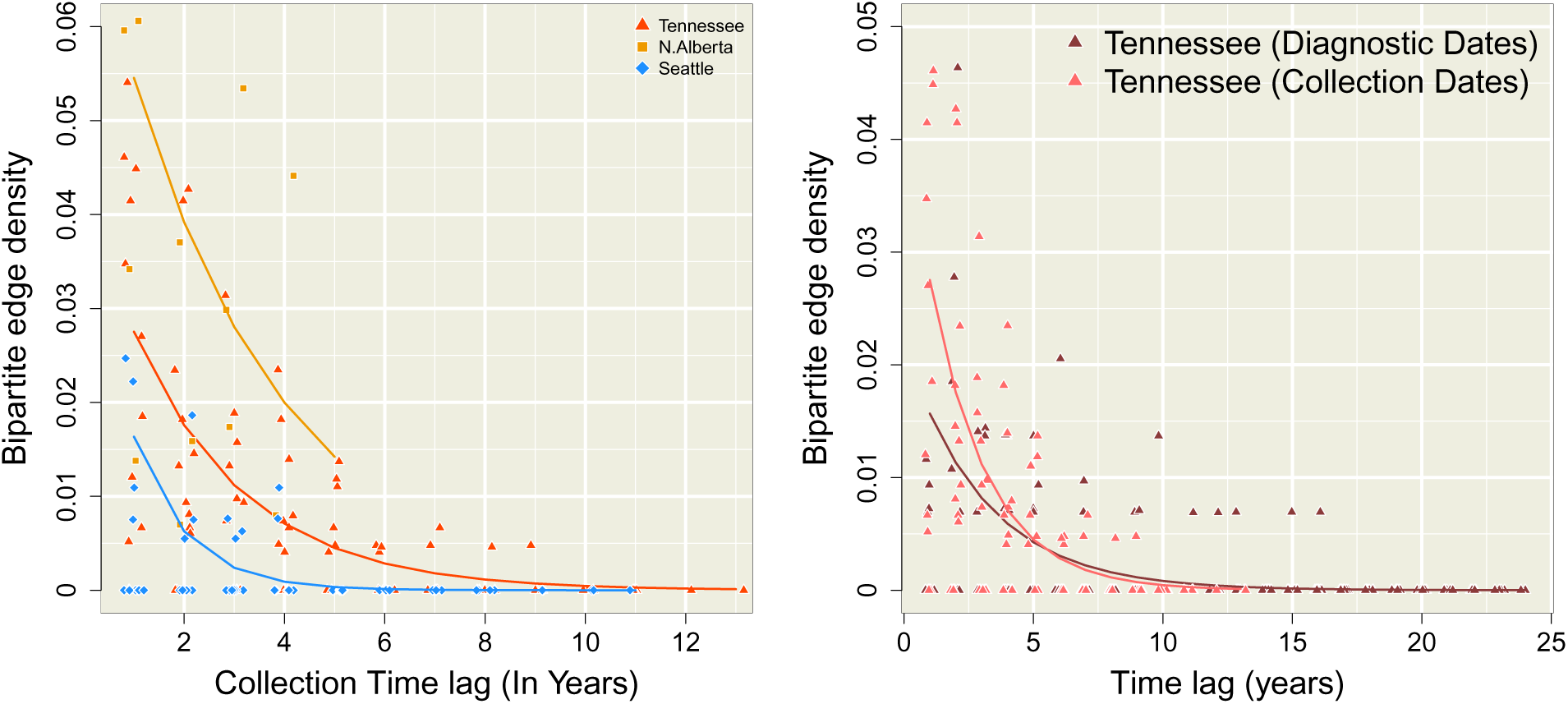
Decay in bipartite edge density with increasing time lag. Each point represents a bipartite graph at a given time lag (difference in years between time points, *x*-axis). We added random noise to the time lag associated with each point to make it easier to distinguish overlapping points. The *y*-axis represents bipartite edge density, *i.e.*, the frequency of edges between cases in different time points out of all edges below *w_max_* = 0.004 (Equation 3). Trend lines represent the predicted edge densities from the binomial regression described by Equation 4. **(left)** The three complete data sets using collection dates. **(right)** The Tennessee data sets filtered for direct comparison between diagnostic and collection date. The diagnostic dates have been corrected by the removal of outlier years from 1982, 1984, and 1986.

**Figure S3:**
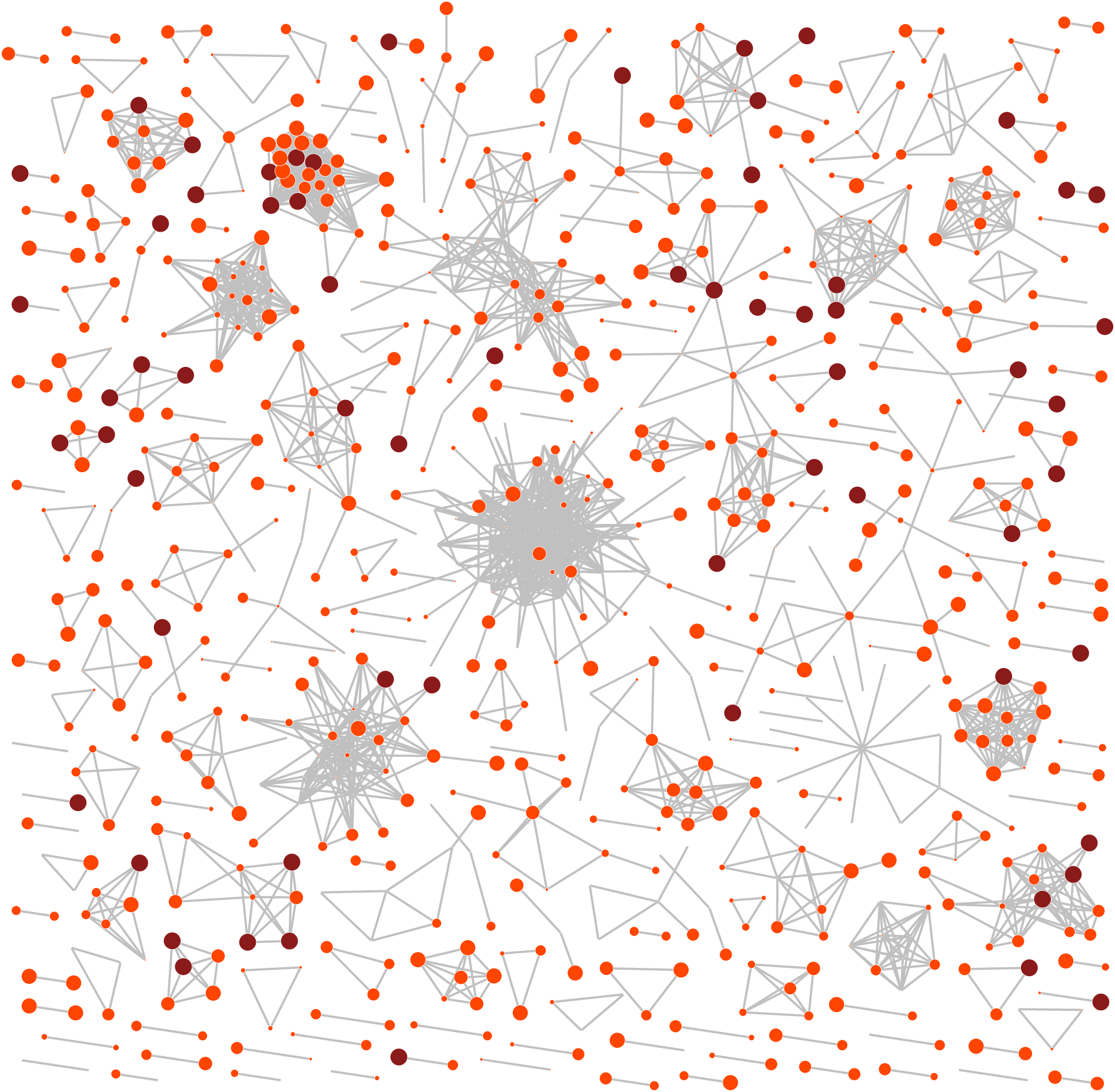
Graph depicting clusters at the GAIC-optimized distance threshold for the Middle Tennessee data set stratified by diagnosis dates instead of sample collection dates. Each point represents a known case (lighter red) or new case (darker red) with sizes scaled in proportion to the year of diagnosis.

**Figure S4:**
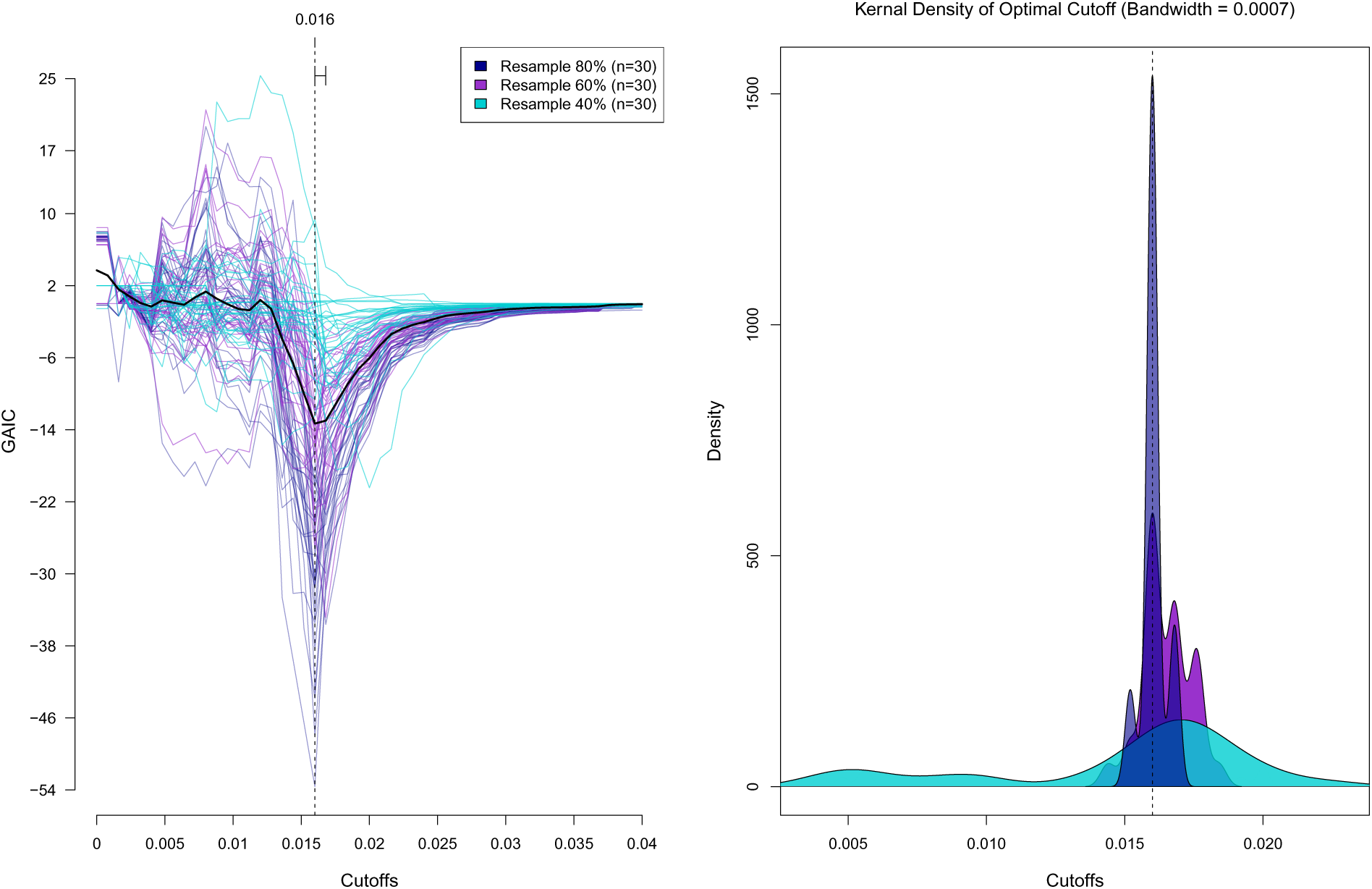
**(left)** GAIC calculated from 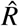 and 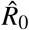 over 51 different cutoffs for 90 random subsamples of the Seattle dataset. The interquartile range (0.0160,0.0168) is indicated at the top of the plot and the 3 different resampling proportions are designated by colour. The solid black line represents a smoothing spline function on the combination of all data sets with an optimal cutoff outlined by the vertical dashed line. **(right)** Gaussian kernal densities of optimal cutoffs (*w*_max_) for each of the three resampling proportions. Each curve represents a sample of *n* = 30 replicate samples.

**Figure S5:**
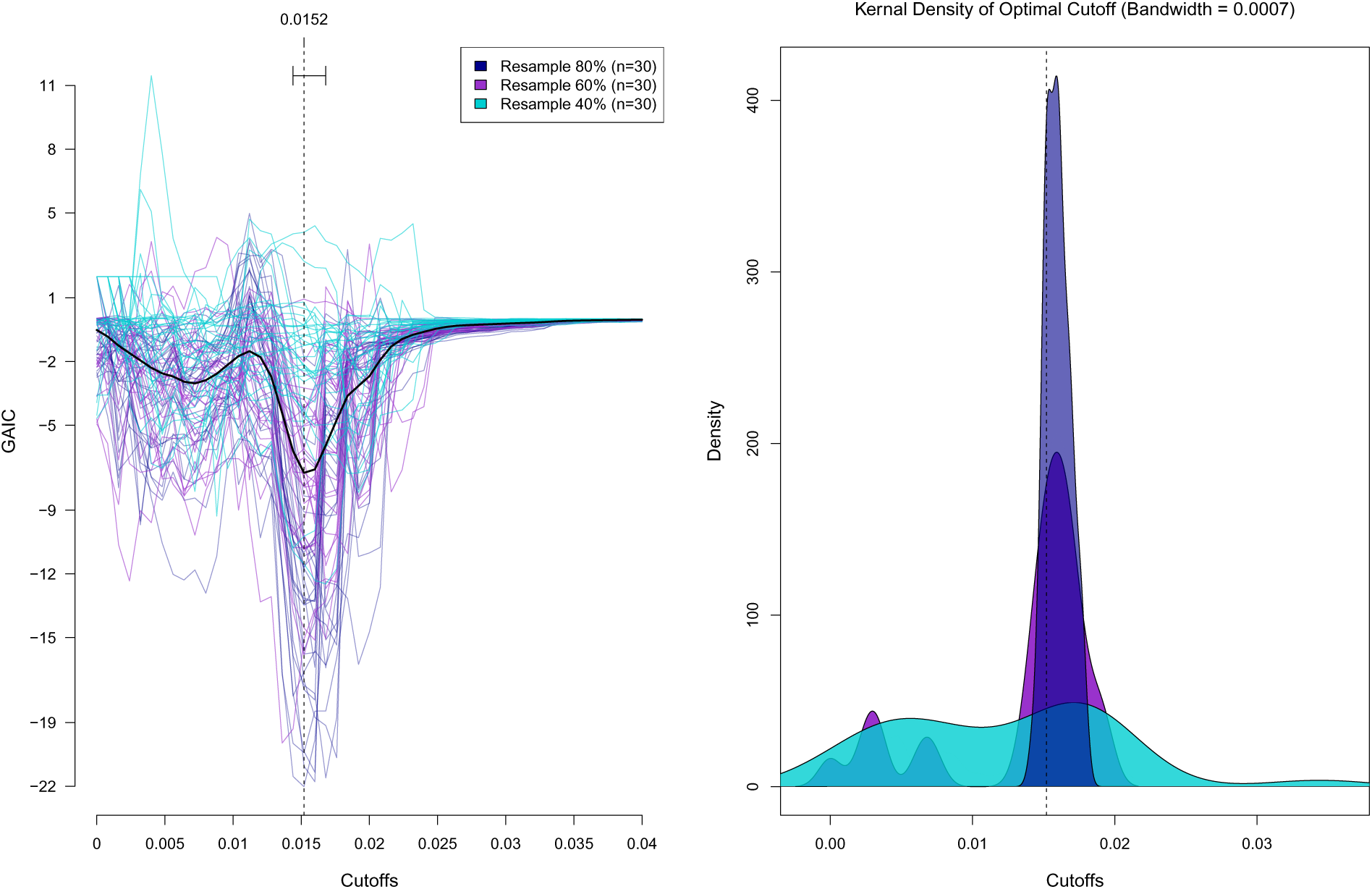
**(left)** GAIC calculated from 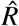 and 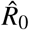 over 51 different cutoffs for 90 random subsamples of the Seattle dataset. The interquartile range (0.0144, 0.0168) is indicated at the top of the plot and the 3 different resampling proportions are designated by colour. The solid black line represents a smoothing spline function on the combination of all data sets with an optimal cutoff outlined by the vertical dashed line. **(right)** Gaussian kernal densities of optimal cutoffs (*w*_max_) for each of the three resampling proportions. Each curve represents a sample of *n* = 30 replicate samples.

**Figure S6:**
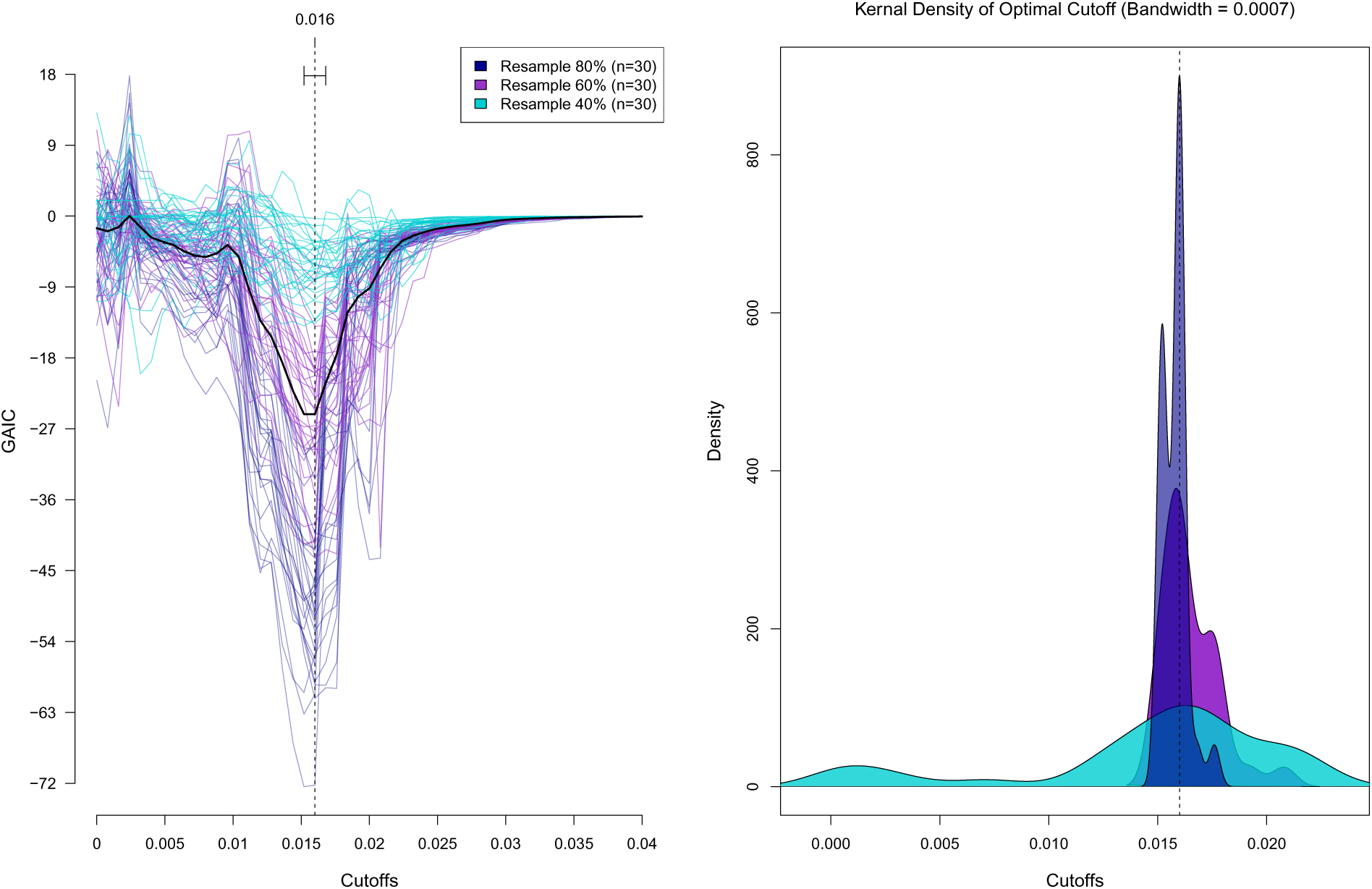
**(left)** GAIC calculated from 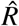 and 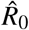 over 51 different cutoffs for 90 random subsamples of the Seattle dataset. The interquartile range (0.0152, 0.0168) is indicated at the top of the plot and the 3 different resampling proportions are designated by colour. The solid black line represents a smoothing spline function on the combination of all data sets with an optimal cutoff outlined by the vertical dashed line. **(right)** Gaussian kernal densities of optimal cutoffs (*w*_max_) for each of the three resampling proportions. Each curve represents a sample of *n* = 30 replicate samples.

**Figure S7:**
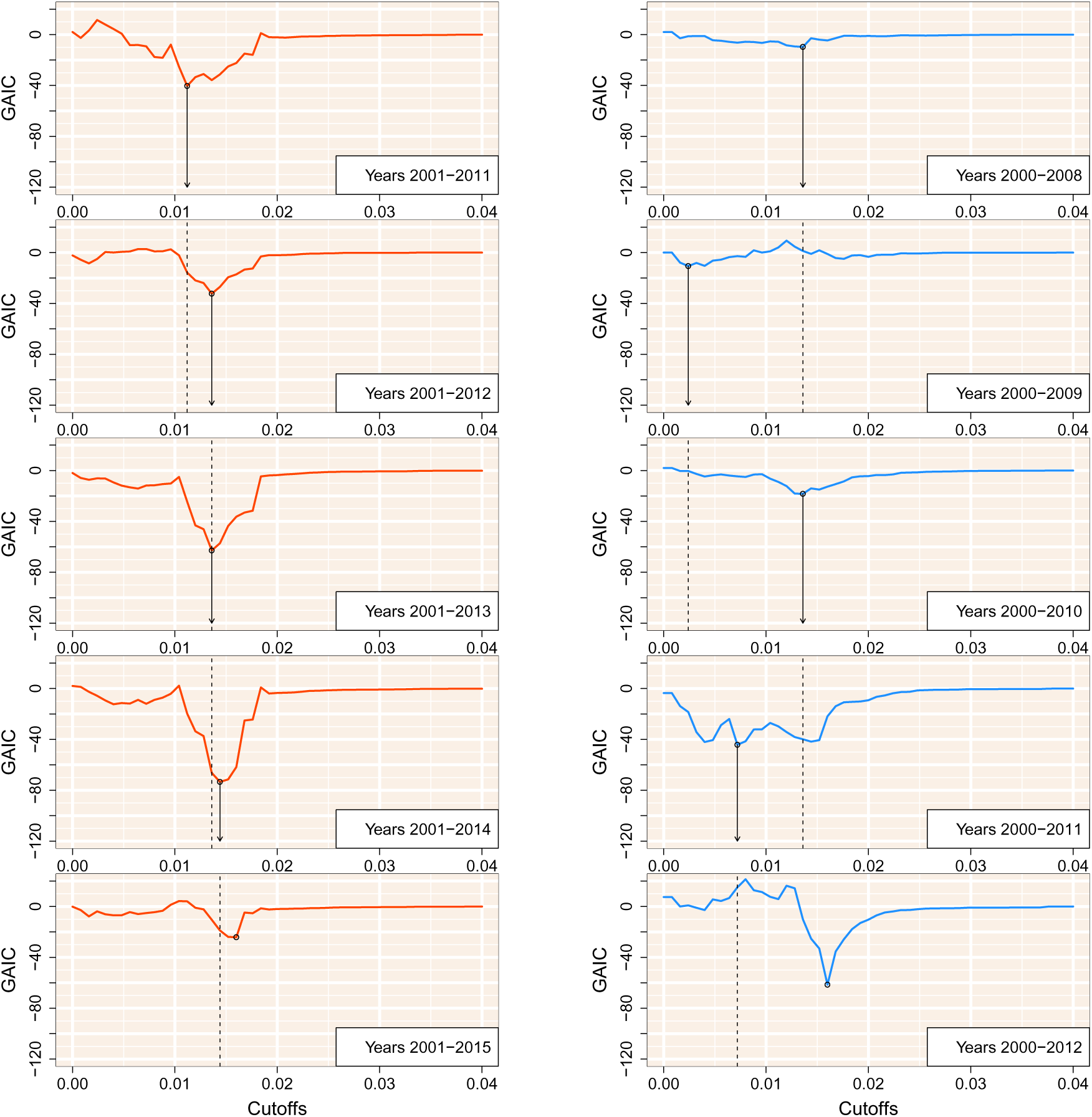
GAIC plotted against the cutoff threshold for 5 sequentially filtered subgraphs of the Complete Tennessee data set (**left**) and the Seattle subset (**right**). The year ranges associated with the GAIC result are shown in the bottom right of each plot and the dashed black line represents the optimum cutoff threshold obtained from the previous result.

## Notes

#### Summary of Updates

Corrections to typographical errors in formulae; mathematical notation changed for clarity; additional data and analyses; figures revised.

